# Exploring the molecular structure of lipids in the design of artificial lipidated antifungal proteins

**DOI:** 10.1101/2024.03.04.583322

**Authors:** Hendra Saputra, Muhammad Safaat, Pugoh Santoso, Rie Wakabayashi, Masahiro Goto, Toki Taira, Noriho Kamiya

## Abstract

Fungal infections have been a concern for decades, yet effective and approved antifungal agents are limited. We recently developed a potential method to enhance the antifungal activity of a small chitin-binding domain (LysM) from Pteris ryukyuensis chitinase A (PrChiA) by the site-specific introduction of a palmitoyl (C16) group catalyzed by microbial transglutaminase (MTG). Herein, we attempted the conjugation of a series of lipid-peptide substrates with LysM genetically fused with a C-terminal MTG-reactive Q-tag (LysM-Q) to yield LysM-lipid conjugates (LysM-lipids) with different lengths (LysM-C12, -C14, and -C16) and different numbers of alkyl chains [LysM-(C12)_2_, - (C14)_2_, and -(C16)_2_]. The enzymatic conjugation proceeded smoothly for all LysM-lipids, except for LysM-(C16)_2_ because of the low aqueous dispersibility of the hydrophobic (C16)_2_ lipid-peptide substrate. The combination of amphotericin B (AmB) with LysM-C14 or LysM-C16 exhibited the highest antifungal performance against Trichoderma viride whereas alterations in the number of alkyl chains were not effective in enhancing the antifungal activity of the LysM-lipids. Fluorescent microscopic analysis showed that the fungal cell wall was stained with C14- and C16-modified LysM-muGFP fusion proteins when combined with AmB, suggesting a synergistic action of AmB and LysM-lipids with a suitable lipid length. All LysM-lipids showed minimum cytotoxicity toward mammalian cells, suggesting that LysM-lipids could be a safe additive in the development of new antifungal formulations.

## Introduction

New antifungal agents that are effective, specific, and safe are required. Drugs with a broad fungicidal activity spectrum are desirable and one of the major broad-spectrum antifungal agents in current use is amphotericin B (AmB)^1^. AmB also affects yeasts and moulds. The mechanism of action of AmB involves making a small hole in the cell membrane through the interaction of AmB with the membrane and the resulting extraction of ergosterol^2,3,4^. The pharmaceutical use of AmB is somewhat restricted because of the high sensitivity to the dosage administered^5^. One potential solution involves reducing the concentrations of AmB to levels that are safer. The use of a mixture of antifungal medications is also a potential alternative^6^ to monotherapy with AmB. Increasing the efficacy of conventional antifungals by chemical modification is another effective strategy. For instance, antifungal properties of gallates in the evaluation against a wood-decaying fungus indicated octyl gallate showed the highest antifungal activity among gallates with different alkyl chain length^7^.

Fungi have cell walls that contain chitin, which is a polysaccharide containing the monomer β-1,4-N-acetyl-glucosamine. Chitin is an important target in the development of antifungal formulations. Chitinase, which is able to degrade chitin into oligomers and/or monomers, causes disruption of the structure of the fungal cell wall, which leads to cell death^8^. Because chitin is absent in humans, targeting chitin in fungal cell walls has potential for the development of safe antifungal agents.

In our previous study, we investigated the potential utility of chitinase A derived from Pteris ryukyuensis (PrChiA)^9^ in antifungal formulations. A potential strategy to enhance the antifungal efficacy of chitinase is via augmenting its binding affinity toward lateral fungal cell walls via hydrophobic interactions^10^. We hypothesized that the use of chemoenzymatic conjugation would be a viable method for increasing the affinity of chitinase toward cell wall components and/or cellular membranes via introducing covalent linkages between lipids and the chitinase proteins^11^. Lipid-conjugated chitinase domains were prepared by a site-specific lipid-protein conjugation catalyzed by microbial transglutaminase (MTG). Interestingly, the combination of AmB solubilized by sodium deoxycholate and artificial palmitoylation (C16) of recombinant PrChiA domains provided a synergistic enhancement of the antifungal activity. In particular, a lysin motif (LysM) found in the carbohydrate-binding domains of PrChiA exhibited higher antifungal activity than the catalytic domain of PrChiA in the growth inhibition of *Trichoderma viride*^12^. In addition, all the palmitoylated chitinase domains tested showed minimal cytotoxicity toward mammalian cells^13^.

The lipid modification of antifungal proteins and enzymes was thus found to be an effective strategy for the development of new antifungal reagents; however, the mechanism of the antifungal action of the palmitoylated LyM domain (LysM-C16) has not been fully elucidated. We have found that the affinity of lipid-protein conjugates with cellular membranes was greatly altered by the structure of lipid moieties^14,15^. The extension of the in vivo half-life of a lipid-modified enhanced green fluorescent protein has been demonstrated through modulation of the affinity with albumin by lipid conjugation^14^. Herein, we explored the effect of the length and the number of alkyl chains of the lipids in the design of artificial lipidated antifungal proteins. Using a LysM domain with an MTG-reactive Q-tag (LysM-Q) as a model, we prepared a series of artificially lipidized LysM (LysM-lipids) (Fig. 1). The results clearly showed that a mono-lipidized with a specific length (C_14_ and C_16_) was effective in enhancing the distribution of the conjugates in fungal cell walls in the presence of AMB solubilized with sodium deoxycholate, implying a synergistic action between the LysM-lipid and AMB. By contrast, introduction of di-lipidized to LysM resulted in aggregate formation, which likely retarded the permeation into the cell walls. The safety of the LysM-lipids was also investigated using mammalian cells.

**Fig. 1.**
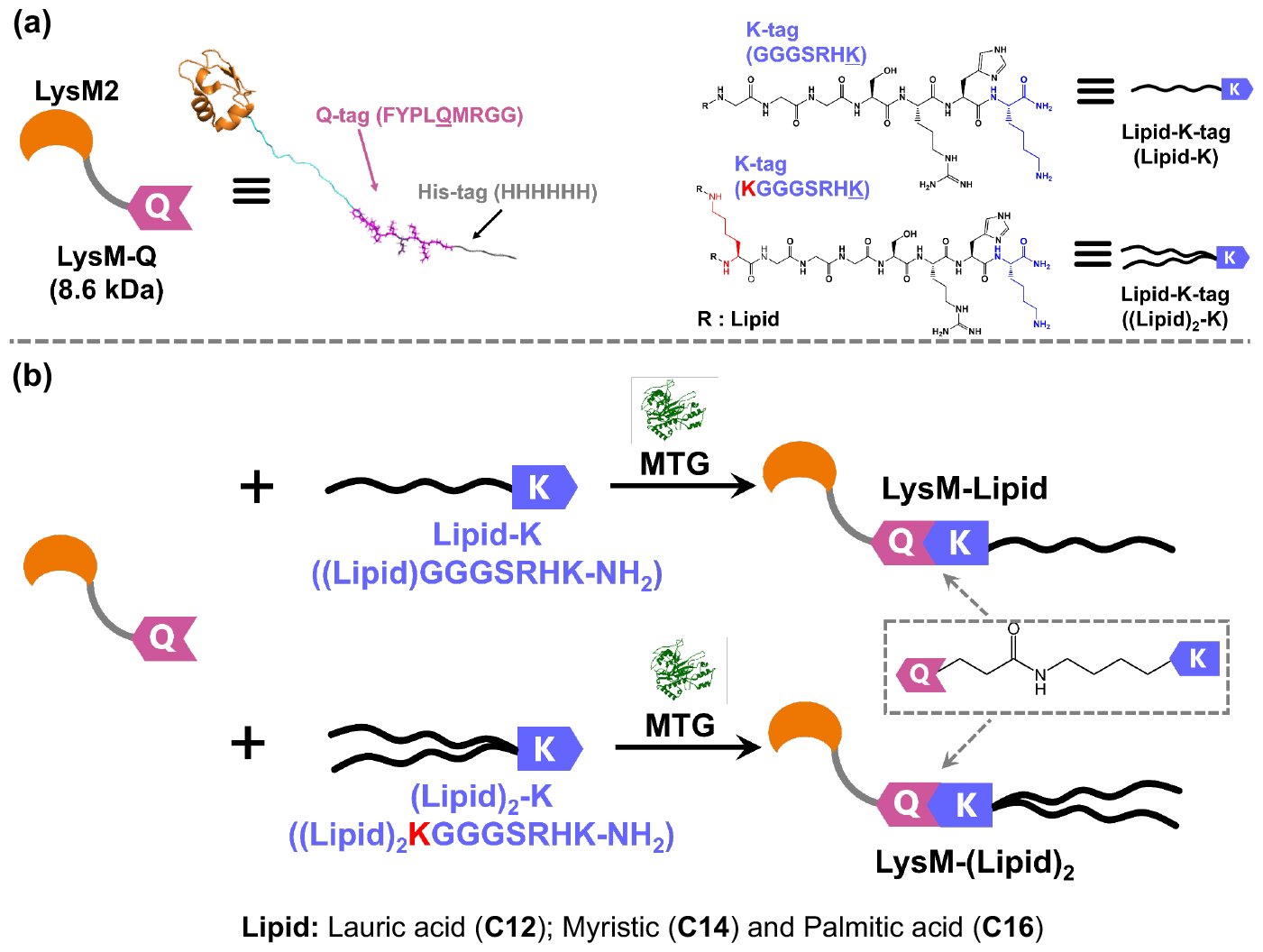
(a) Schematic illustration of the LysM domain derived from PrChiA with a C-terminal Q-tag (LysM-Q). The Q-tag is an MTG-reactive peptide tag containing a glutamine residue (FYPLQMRGG). (b) MTG-catalyzed bioconjugation of Q-tagged LysM and lipid peptide substrates containing an MTG-reactive Lys residue in the peptide moiety (GGGSRHK).

## Results and discussion

### Conjugation of LysM with different types of lipid substrates

MTG-catalyzed crosslinking of LysM-Q with a mono-lipidized peptide substrate containing an MTG-reactive LysM residue (C12-K, C14-K, or C16-K) or with a mono-lipidized peptide substrate [(C12)_2_-K, (C14)_2_-K, or (C16)_2_-K] was conducted to yield LysM-lipid conjugates (Supplementary Fig. S2). The resultant conjugates were designated as LysM-C12, -C14, and -C16 (with mono-lipidized) and LysM-(C12)_2_, -(C14)_2_, and -(C16)_2_ (with mono-lipidized). The results of sodium dodecyl sulfate-polyacrylamide gel electrophoresis (SDS-PAGE) showed a clear band shift between MTG-modified and unmodified LysMs, except for LysM-(C16)_2_. A single product band was observed for each LysM-lipid conjugate after the MTG conjugation reaction, suggesting little formation of multiply labeled byproducts (Supplementary Fig. S3)^12,13^. In the case of LysM-(C16)_2_, the conjugation reaction was incomplete. Because the reaction mixture with (C16)_2_-K was translucent, the MTG-catalyzed conjugation was likely hindered by aggregation of the dipalmitoylated lipid substrate. LysM-(C16)_2_ was thus not included for further assessment.

### Evaluation of antifungal activity

The results of the antifungal activity tests are shown in Figs. 2 and S4. Alone, AMB effectively inhibited fungal growth at 2.5 µM. When LysM-Q (1 µM) was mixed with AMB at 1.25 µM, synergistic inhibition was observed, resulting in the complete suppression of fungal growth (Fig. S4). From a quantitative evaluation (Fig. 2a), LysM-C12 showed a higher level of antifungal activity compared with unmodified LysM (LysM-Q), but lower activity than that of LysM-C14 or LysM-C16. The antifungal activity of LysM-C14 and LysM-C16 at a concentration of 1 μM was similar and complete suppression of the growth of T. viride for 60 h was observed when combined with 0.63 µM AMB. These results were consistent with those of our previous reports; however, we found that not only palmitoylation (LysM-C16) but also the myristoylation of LysM (LysM-C14) was effective in enhancing the antifungal activity of LysM. In addition, the introduction of a lauroyl moiety (LysM-C12) did not achieve antifungal activity comparable to LysM-C14 or LysM-C16.

**Fig. 2.**
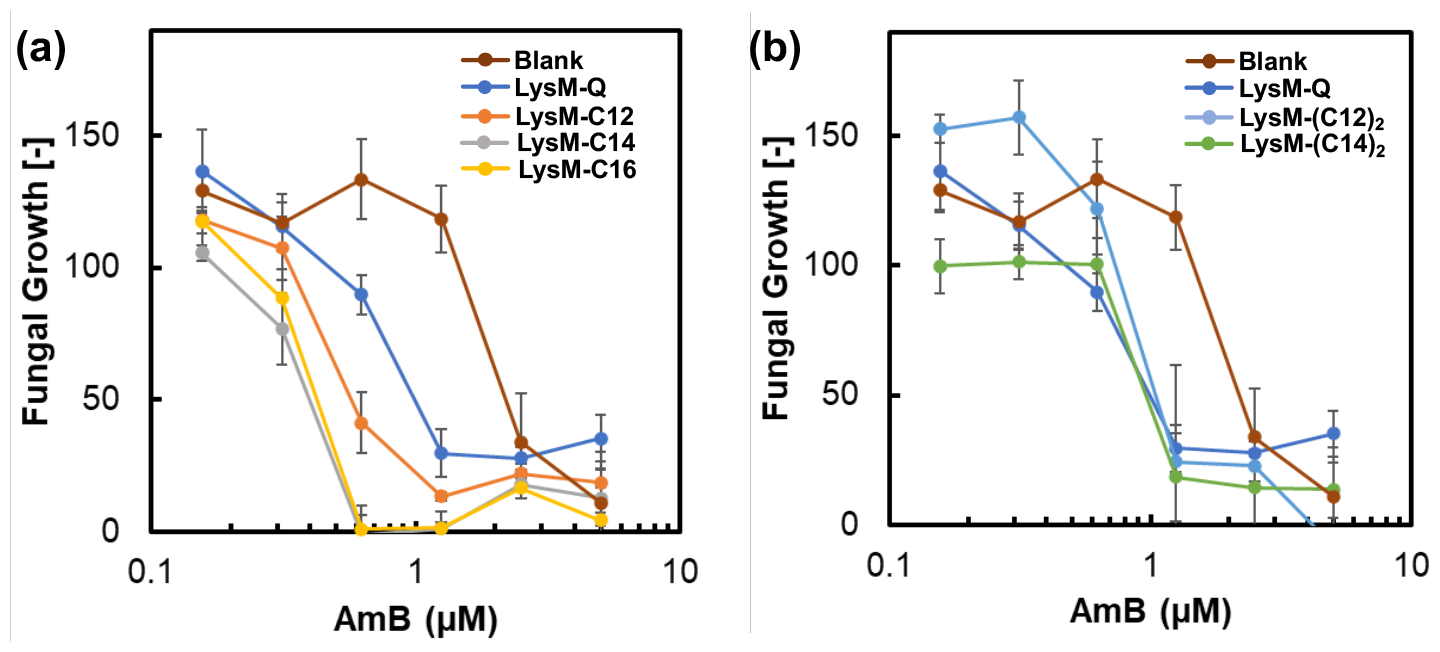
Enzymatic activity assay of K10X/Y12A MTGz mutants by a hydroxamate assay. The underlined K denotes the Y12A single mutant (n.d.: Not detected, below the detection limit). Data are presented as the mean ± standard deviation of three experiments (*n* = 3).

Next, we evaluated the effect of dilipidation of LysM on the antifungal activity. Unexpectedly, both LysM-(C12)_2_ and LysM-(C14)_2_ exhibited lower levels of antifungal activity compared with LysM modified with mono-lipidized and the same level of antifungal activity as unmodified LysM (Fig. 2b). The antifungal activity of all the tested dilipidated LysMs was independent of the length of the alkyl chains. An increase in the number of alkyl chains generally increases the hydrophobicity, which would enhance the affinity with cellular membranes. However, an increase in the hydrophobicity caused by an increase in the length and number of the alkyl chains can decrease the dispersibility in aqueous solutions. In the case of our artificial lipid-modified antifungal proteins, the latter case seemed predominant, leading to there being no significant change in activity between unmodified and dilipid-modified LysMs.

Our previous studies using enhanced green fluorescent protein (EGFP) have shown the impact of the lipid alkyl chain length on the anchoring ability to the plasma membrane of mammalian cells^16^. The cell anchoring ability of EGFP-C12, -C14, and -C16 was associated with the length of the alkyl chain and there was a clear difference between EGFP-C14 and EGFP-C16. We hypothesized that the antifungal action of LysM-lipids would also increase with increasing alkyl chain length because of an increased ability for anchoring to lipid bilayers. The similar antifungal activity of LysM-C14 and LysM-C16 suggested that the antifungal action of LysM-lipids cannot be explained solely by the cell membrane-anchoring ability and likely reflects the permeability into the fungal cell wall. To better understand the antifungal behavior of di-lipidated LysMs, we evaluated the aggregation of LysM-lipids in aqueous solution.

### Dynamic light scattering (DLS) analysis of LysM-lipids

To gain more information on the state of LysM-lipids in an aqueous solution with and without AMB, we investigated the size distribution of LysM-lipids by DLS measurements. The particle sizes and polydispersity index (PDI) values of the different LysM-lipids are listed in Table 1 and the particle size distributions are shown in Supplementary Fig. S5. The results showed that lipid modification led to an increase in the average size of the aggregates compared with the unmodified LysM-Q. As the length of the alkyl group increased, a corresponding increase in the size of the aggregates was observed. An increase in the number of alkyl chains also resulted in an increase in the size of the aggregates. These results suggested that LysM-lipids self-assembled in aqueous solution.

**Table 1.** DLS measurements of AmB with LysM-Q (1 µM) or LysM-lipid (1 µM) in 20 mM NaPi, pH 7.4 at 25 °C. The error bars represent the standard error of three individual measurements. PDI: polydispersity index. The concentrations of AMB were set to those that could suppress fungal growth based on the results shown in Fig. 2.

| # | Samples | Size (nm) |  | PDI |  |
| --- | --- | --- | --- | --- | --- |
|  |  | w/o AmB | with AmB | w/o AmB | with AmB |
| 1 | LysM-Q | 5.0 ± 1.4 | 18.2 ± 0<sup>§</sup> | 0.614 ± 0.013 | 0.276 ± 0.104<sup>§</sup> |
| 2 | LysM-C12 | 16.6 ± 2.7 | 20.2 ± 3.6<sup>§§</sup> | 0.718 ± 0.013 | 0.396 ± 0.081<sup>§§</sup> |
| 3 | LysM-(C12)<sub>2</sub> | 21.2 ± 3.1 | 23.3 ± 1.9<sup>§</sup> | 0.390 ± 0.021 | 0.414 ± 0.029<sup>§</sup> |
| 4 | LysM-C14 | 17.5 ± 3.1 | 21.2 ± 3.1<sup>§§</sup> | 0.432 ± 0.072 | 0.492 ± 0.101<sup>§§</sup> |
| 5 | LysM-(C14)<sub>2</sub> | 22.2 ± 1.9 | 27.1 ± 4.8<sup>§</sup> | 0.321 ± 0.082 | 0.381 ± 0.021<sup>§</sup> |
| 6 | LysM-C16 | 18.2 ± 0.0 | 21.0 ± 0<sup>§§</sup> | 0.475 ± 0.013 | 0.474 ± 0.013<sup>§§</sup> |

<sup>§</sup> AmB [1.25 μM]; <sup>§§</sup> AmB [0.63 μM]

All formulations of LysM-Q or mono-lipidated LysMs exhibited an increase in size upon mixing with AMB. However, LysM-(C12)_2_ and LysM-(C14)_2_ with AMB showed little change in the size distribution before and after the addition of AMB (Fig. S5). In addition, the PDI values showed little variation upon combination with AMB. It is noted that a commercial formulation of AMB contains sodium deoxycholate to solubilize AmB. Therefore, the change in the size distribution of mono-lipidated LysMs upon mixing with AMB may reflect different aggregation states in the solution^17^. This may be because the hydrophobic AMB molecules are associated with the lipid moieties of LysM-lipids and this affects the structure of the micelles. The results also suggested that the monolipidated LysMs can easily form mixed aggregates with AmB and sodium deoxycholate, which will lead to co-permeation of the three components into the fungal cell wall. By contrast, the slight change in the size distribution of mono-lipidized LysM-lipids in the presence of AMB suggested the formation of more stable aggregates because of the increase in the hydrophobic moieties. The formation of stable aggregates by LysM-(C12)_2_ and LysM-(C14)_2_ may retard the penetration into the fungal cell membrane.

### Distribution analysis of LysM-muGFP-lipids using confocal laser scanning microscopy (CLSM)

Further investigations were carried out using a fusion protein of LysM and green fluorescent protein (muGFP) with a *C*-terminal MTG-reactive Q-tag (LysM-muGFP-Q)^13^ to study how the formulation interacted with *T. viride* (Figs. 3 and S6). The blue fluorescence in the control group showed the presence of chitin that constitutes the fungal cell wall. LysM-muGFP-Q combined with AMB showed a weak overlap of green (from muGFP) and blue (from CFW, calcofluor white, as a staining agent for chitin determination) fluorescence. The behavior of LysM-muGFP-C12 was similar to that of LysM-muGFP-Q, which indicated that the effect of the introduction of the C12 lipid was not significant, yet both proteins could bind to chitin in the fungal cell wall. By contrast, marked accumulation of the green fluorescence derived from LysM-muGFP-C14 and LysM-muGFP-C16 was observed with AMB compared with the other formulations. The presence of the mono-lipidized C12 or C14 with LysM-muGFP-(C12)_2_ or LysM-muGFP-(C14)_2_ significantly reduced the green fluorescence in the cell wall. Overall, the trends for the green fluorescence intensity derived from LysM-muGFP-lipids in the fungal cell wall (Fig. 3) were in good agreement with the results observed for antifungal activity (Fig. 2).

**Fig. 3.**
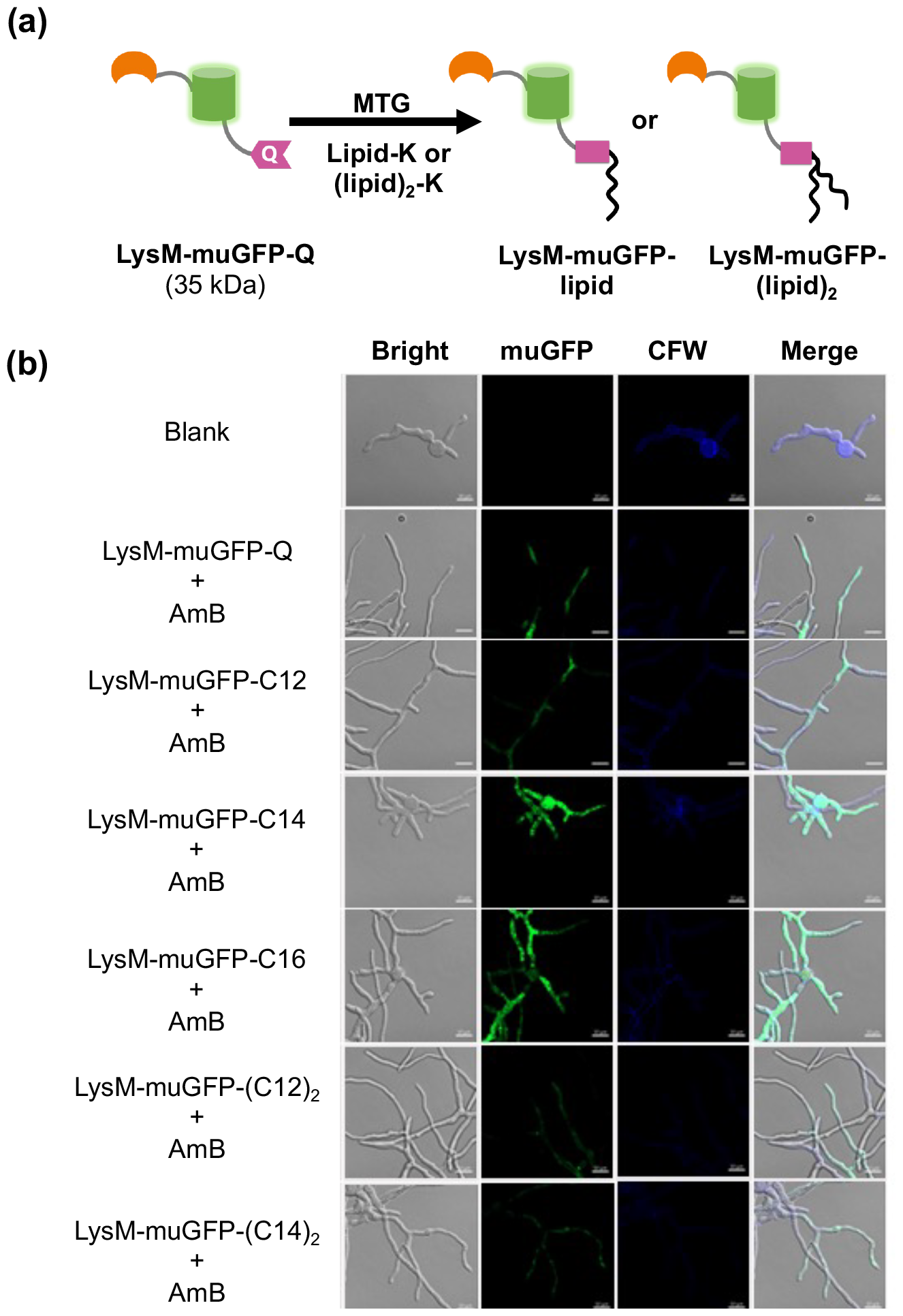
(a) Schematic illustrations of the bio-conjugation between LysM-muGFP-Q with lipid-K or (lipid)2-K catalyzed by MTG. (b) CLSM analysis of AmB with LysM-muGFP-Q or LysM-muGFP-lipid(s) in the presence of T. viride hyphae in 20 mM NaPi, pH 7.4, at 25°C (bars: 10 μm).

Our previous results for a liposomal formulation of AmB (AmBisome®) showed that the accumulation of LysM-muGFP-C16 occurred primarily at the tip of the hypha^13^. In the present study, the whole fungal body was fluorescently stained with LysM-muGFP-C14 and LysM-muGFP-C16 in the presence of AMB, but there was no significant staining in the absence of AMB (Fig. S6). The strong green fluorescence observed with LysM-muGFP-C14 and LysM-muGFP-C16 clearly indicated that these recombinant proteins were captured on the fungi by the interaction of the LysM moiety with the fungal cell wall chitin. These results indicated that LysM modified with a single, hydrophobic alkyl chain nonspecifically attached to the fungal cell wall chitin in association with the action of AmB solubilized by sodium deoxycholate.

The treatment of fungi with LysM-muGFP-(lipid)_2_ showed a notable pattern that differed from that observed for the mono-lipidized LysM-muGFP-lipids. When combined with AMB, little alteration in the average size of the aggregates was observed for LysM-muGFP-(lipid)_2_ (Fig. S5). When mixed with α-chitin, green fluorescence was observed in the CLSM analyses for all the samples [LysM-muGFP-Q, LysM-muGFP-lipid, and LysM-muGFP-(lipid)_2_] (Fig. S7), suggesting that the free LysM moiety and LysM in each lipid conjugate retained the binding ability to chitin. However, the impact of hydrophobicity appeared pronounced with an increase in the number of alkyl chains. We observed small dots with green fluorescence on α-chitin for LysM-(C12)_2_ and LysM-(C14)_2_, implying the formation of more stable aggregates with LysM-muGFP-(lipid)_2_ than with LysM-muGFP-lipid. This stable aggregation may lead to a reduction in the antifungal efficacy.

Collectively, these results indicated that LysM-lipids and AmB appear to have a specific mode of antifungal action. The process may begin with the action of AmB, which has been shown to bind to ergosterol in the cell membrane and subsequently form holes^18^. This hole formation is then followed by the action of LysM-lipids with sufficient hydrophobicity (C14 and C16), which synergistically enhanced the detrimental effects of AmB on the fungal cell wall. However, increasing the hydrophobicity over an optimum value decreased the antifungal activity possibly because of the strong self-association of dilipid-modified LysMs. Increasing the hydrophobicity could increase the hemolytic activity of formulations^19^, and thus we tested the cytotoxicity of all the LysM-lipids.

### Cytotoxicity testing

The cytotoxicity of the LysM-lipids was evaluated (Fig. 4). The LysM-lipids showed no detrimental effects on Hek293T cells and these results were consistent with our previous studies with LysM-C16^12,20^. The treatment of cells with 1 µM LysM-lipid, and ≤1.25 µM LysM-lipid in the presence of AMB, showed little effect on the cell viability. In the presence of AMB, the cell viability was slightly decreased to ca. 87% possibly because of the effect of the co-surfactant, but no significant difference was observed by altering the length or the number of the alkyl chain moieties of the LysM-lipids.

**Fig. 4.**
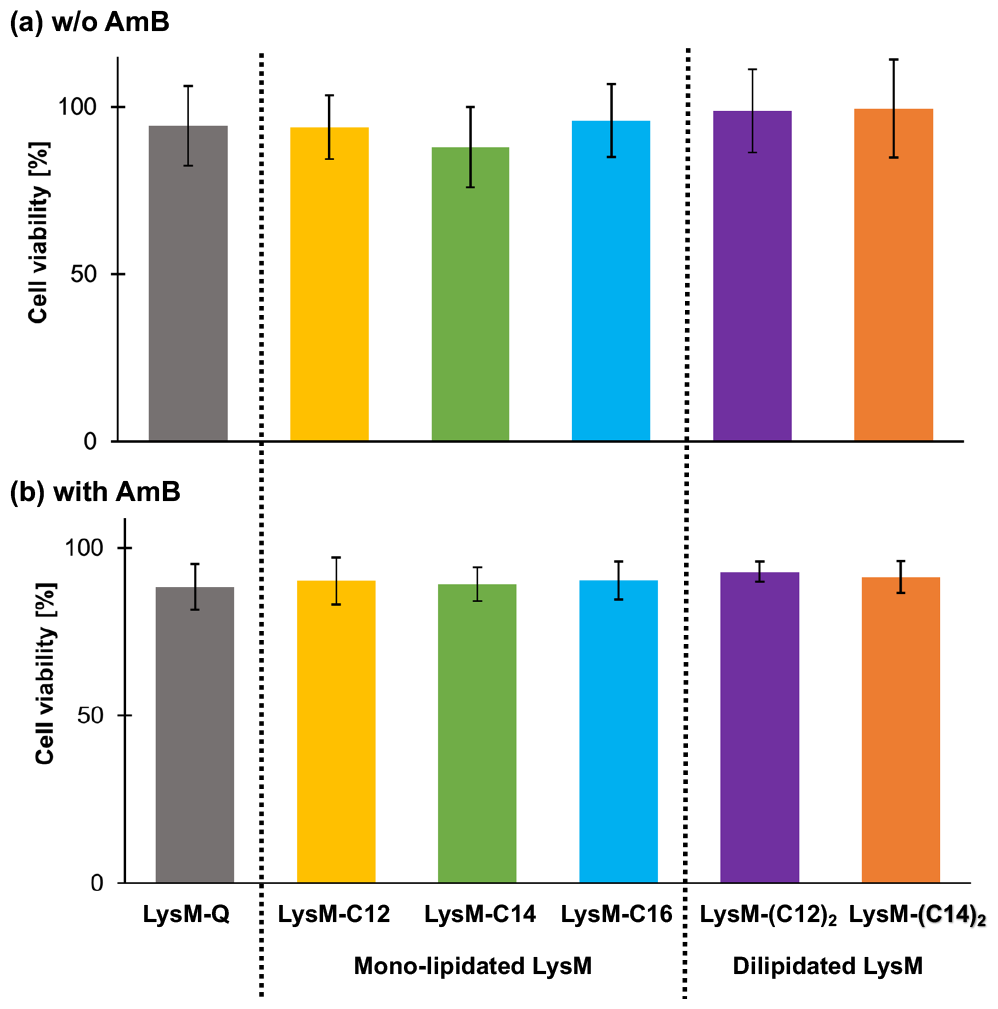
Cell viability of Hek293T cells (5,000 cells/well) incubated with LysM-lipids alone (a) and in combination of AmB (b). Cell viability was measured using a Cell Counting Kit-8 (Dojindo).

Cytotoxicity testing was also conducted using HeLa cells (a cancer cell line). These results also showed little negative effect on the cell viability (Supplementary Fig. S8). The effect of the alkyl chain length on the cytotoxicity properties has been previously evaluated in SNU-1 cells, which resulted in findings consistent with the present study showing little cytotoxicity of EGFP-lipids. It was also indicated that β-sheet formation of the lipid-peptide substrates did not damage the cell membrane significantly, but the hydrophobicity of the lipid moiety determined the critical micelle concentration and cytotoxicity^14^. These results showed that LysM-lipids would be safe to use as an additive for the development of new antifungal formulations.

## Conclusion

The present study aimed to assess the impact of the length and the number of alkyl chains on the antifungal activity of site-specific, lipid-modified LysMs prepared using MTG. The cross-linking efficiency was independent of the type of the lipid-peptide substrate, and the conjugation proceeded quantitatively, except for (C16)_2_-K, which showed less aqueous dispersibility compared with the other lipid-peptide substrates. The antifungal activity of the LysM-lipids was influenced by both the length and the number of the alkyl groups, and specifically LysM-C14 and LysM-C16 showed the highest antifungal activity against *T. viride*. By contrast, increasing the number of C12 or C14 alkyl chains markedly reduced the antifungal activity, possibly because of the formation of tight aggregates compared with those formed from mono-lipidized LysM-lipids. These data will provide new insights into the development of antifungal strategies that target plasma membrane components of pathogenic fungi, such as ion channels and G protein-coupled receptors^21^.

## Materials and methods

### 1. Materials

Luria−Bertani (LB) broth medium, ammonium peroxodisulfate, 30% acrylamide/bis mixed solution (29:1), tris (hydroxymethyl)aminomethane, tryptone, dried yeast extract, dipotassium hydrogen phosphate, and hydrochloric acid were purchased from Nacalai Tesque, Inc. (Kyoto, Japan). Sodium dodecyl sulfate (SDS), glycerol, potassium dihydrogen phosphate, and sodium chloride were purchased from Wako Pure Chemical Industries, Ltd. (Osaka, Japan). *N,N,N′,N′*-Tetramethylethylenediamine, HisTrap FF crude 5 mL column, HiTrap Q HP column 5 mL, PD SpinTrap G-25, and Ni Sepharose 6 Fast Flow were purchased from Cytiva (Tokyo, Japan). Amicon Ultra-0.5 (PLBC Ultracel-3 membrane, 3 kDa) and Amicon® Ultra-15 Centrifugal Filters (3 kDa MWCO) were from Millipore (Tokyo, Japan). Calcofluor-white and imidazole were from Sigma-Aldrich (Tokyo, Japan). Gibco AMB containing 250 μg of AMB and 205 μg of sodium deoxycholate was purchased from ThermoFisher Scientific. An α-chitin nanofiber (α-CNF) provided by professor Shinsuke Ifuku (Tottori University) was used for the chitin binding assay. The width and degree of deacetylation of α-CNF prepared from crab shell powder is 10–20 nm and 3.9%, respectively^22^.

### 2. Expression and Purification of Recombinant Proteins

The preparation and use of the proteins MTG, LysM-Q, and LysM-muGFP-Q were performed as described previously^13,23^.

### 3. Fmoc Solid Phase Peptide Synthesis

The lipid-peptide substrates, lipid-G_3_S-RHK-NH_2_and (lipid)_2_-KG_3_S-RHK-NH_2_, were produced manually using established Fmoc solid phase peptide synthesis methodology, in accordance with previous studies^14^.

### 4. MTG-catalyzed Lipid-conjugation Reaction

Briefly, the bioconjugation of LysM-Q with mono-lipidized (C12-K, C14-K, and C16-K) and mono-lipidized [(C12)_2_-K, (C14)_2_-K, and (C16)_2_-K] lipids was performed in a total reaction volume of 500 µL. The reaction components (100 µM C14, 10 µM LysM-Q, 1% n-dodecyl-β-D-maltoside (DDM), and 0.1 U/mL MTG) were dissolved in 10 mM Tris HCl buffer pH 7.4 and reacted in an incubator shaker. The reaction was performed at 180 rpm and 37°C for 1 h. Purification was carried out with Ni-Sepharose 6 Fast Flow resin, then using a PD SpinTrap G25 column to obtain the conjugated protein, and finally the sample was concentrated.

### 5. Chitinase-AmB Formulation Antifungal Activity Testing

AmB (0−5 μM), protein, lipid (1 μM), and *T. viride* (10,000 spores/mL) were mixed in potato dextrose broth (PDB) in 60-μL 96-well plates at 25°C and 85% humidity. *T. viride* mycelial growth at 60 h was examined by measuring the optical density (OD) values of the growing fungus using ImageJ to determine well intensity^12^.

### 6. Particle size analysis using DLS

The particle size of the LysM-lipid and AmB mixture in 20 mM sodium phosphate butter (NaPi) (pH 7.4) with a total volume of 100 μL was determined using DLS.

### 7. Confocal Laser Scanning Microscopy (CLSM) analysis

CLSM was performed according to a previous report^13^. In brief, 1 × 10^6^ *T. viride* spores were added to 10 mL of autoclaved PDB medium. The mixture was then kept at 25°C and shaken at 150 rpm for 18 h to obtain *T. viride* mycelia. The mycelial suspension underwent centrifugation at 3000 × g and 25°C for 20 min to gather the mycelia. Then, a small quantity of PDB medium was reintroduced to re-suspend the mycelia. Each sample was prepared with the optimal concentration for the antifungal testing and Calcofluor-white (CFW) staining. CFW is a non-specific fluorochrome that binds with cellulose and chitin in the cell walls of fungi and other organisms. Subsequently, the sample was incubated at 25°C for 1 h. The samples underwent a triple washing procedure using a solution of 20 mM NaPi (pH 7.4) before being examined using CLSM (LSM700; Carl Zeiss, Oberkochen, Germany).

### 8. Cytotoxicity testing

A total of 5 × 10^3^ Hek293T and HeLa cells were put into individual wells containing Dulbecco’s Modified Eagle Medium (D-MEM) [containing 10% Fetal Bovine Serum (FBS) and 1% Antibiotic-Antimycotic solution (Gibco)]. The cells were then left to grow for 24 h at 37°C and 5% CO_2_. After letting the plates sit for 24 h, samples containing the correct amount of each formulation determined according to an effective fungal growth inhibition were put into each well. A Cell Counting Kit-8 (CCK-8; Dojindo, Kumamoto, Japan) was used to determine the cell viability. The wells were treated for 3 h with 10 µL of CCK-8 solution. Then, the absorbance at 450 nm was measured using a microplate reader. The following formula, where A_test_, A_blank_, and A_control_ are the absorbances of the test, blank, and control samples, respectively, was used to determine the cell viability^13^.

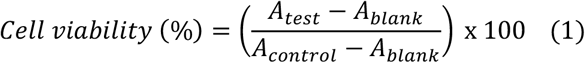

## Acknowledgments

This study was supported by JSPS KAKENHI Grant numbers JP19H00841 and JP23H00247 (to N.K.). We greatly appreciate Prof. Shinsuke Ifuku (Tottori University) for providing α-chitin nanofiber. All authors have provided consent.The authors thank Mr. Kazuki Uchida (Kyushu University) for the assistance in experiments and valuable discussion. We thank Victoria Muir, PhD, from Edanz (https://jp.edanz.com/ac) for editing a draft of this manuscript.

