## Supplemental figures for "Exploring the molecular structure of lipids in the design of artificial lipidated antifungal proteins"

### Supporting information

#### 1-1. Amino acid sequence of LysM-Q (pI/Mw: 6.99 / 8739.65)

MCTTYTIKSGDTCYAISQARGISLSDFESWNAGIDCNNLQIGQVVCVSKPSTSTTPSPTPSSSSN
GFYPLQMRGGHHHHHH

#### 1-2. Amino acid sequence of LysM-muGFP-Q (pI/Mw: 6.04 / 35078.23)

MCTTYTIKSGDTCYAISQARGISLSDFESWNAGIDCNNLQIGQVVCVSKPSTSTTPSPTPSSSSN
GHHHHHHSKGEELFTGVVPILVELDGDVNGHKFSVRGEGEGDATNGKLTLKFICTTGKLPVP
WPTLVTTLTYGVLCFSRYPDHMKRHDFFKSAMPEGYVQERTISFKDDGTYKTRAEVKFEGDT
LVNRIELKGIDFKEDGNILGHKLEYNFNSHNVYITADKQKNGIKAYFKIRHNVEDGSVQLADH
YQQNTPIGDGPVLLPDNHYLSTQSVLSKDPNEKRDHMVLLEDVTAAGITHGMDELYRGGGGS
LLQG

Brown: LysM2 domain; Blue: Linker sequences derived from PrChiA; Green: Hexahistidine tag;
Purple: Q-tag (FQ); Dark Green: muGFP; Yellow: Q-tag (LQ)

#### 1-3. Molecular structure of lipid-modified peptides

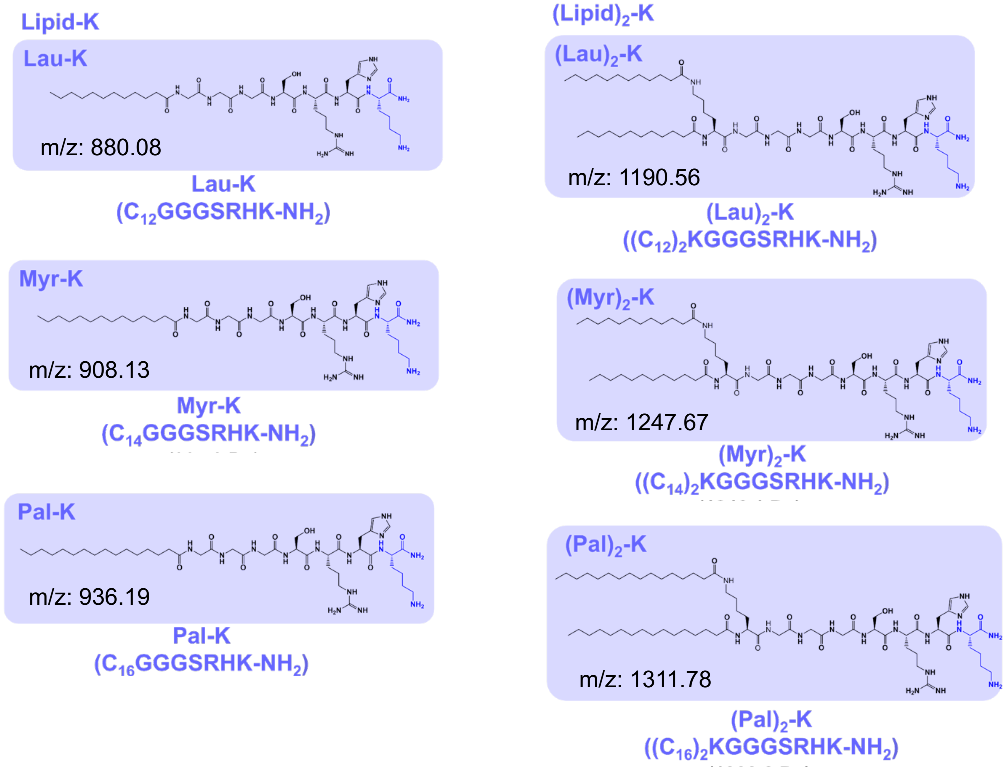


**Fig. S1** Chemical structures of lipid-$G_3$S-RHK and (lipid)$_2$-K$G_3$S-RHK with the values of theoretical molecular weight.

### 2. Supplementary results

#### 2-1. Results of Fmoc solid phase peptide synthesis

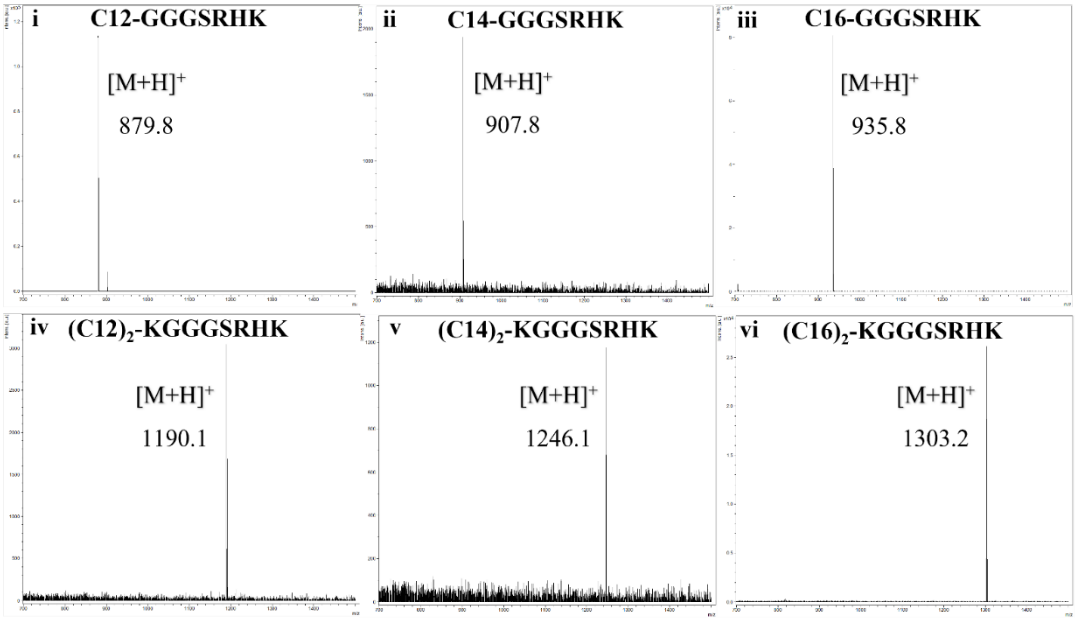


**Fig. S2** MALDI-TOF-MS results of lipid-GGGS-RHK and (lipid)$_2$-KGGGS-RHK. (**i**) C12-, (**ii**) C14-, (**iii**) C16-GGGS-RHK, (**iv**) (C12)$_2$-, (**v**) (C14)$_2$-, and (**vi**) (C16)$_2$-KGGGS-RHK.

#### 2-2. Conjugation of Q-Tagged chitinase domains by MTG

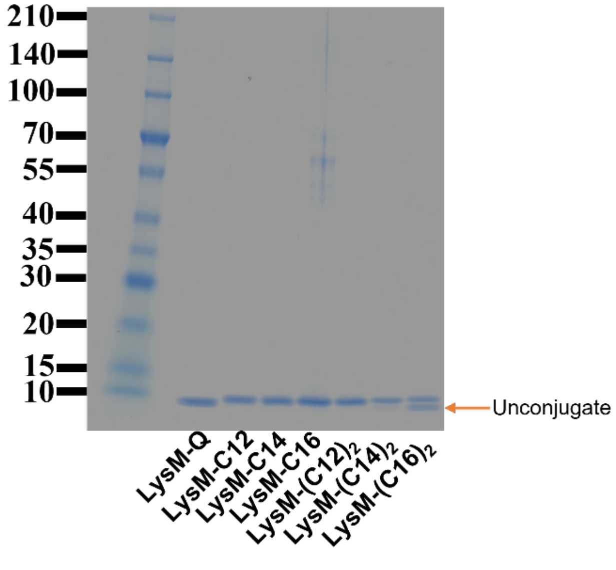


**Fig. S3.** Bioconjugate of the Q-tagged chitinase domains. (SDS-PAGE analysis results of unmodified chitinase and chitinase modified with C12-K, C14-K, C16-K, (C12)$_2$-K, (C14)$_2$-K and (C16)$_2$-K by MTG. All conjugation reactions were carried out under conditions of 10 μM Q-tagged chitinase domains, 1% DDM, 10 μM Lipid-K, and 0.1 U/mL MTG in 10 mM Tris-HCl (pH 8.0) at 37 °C for 1 h.

### 2-3. Qualitative results of antifungal activity test

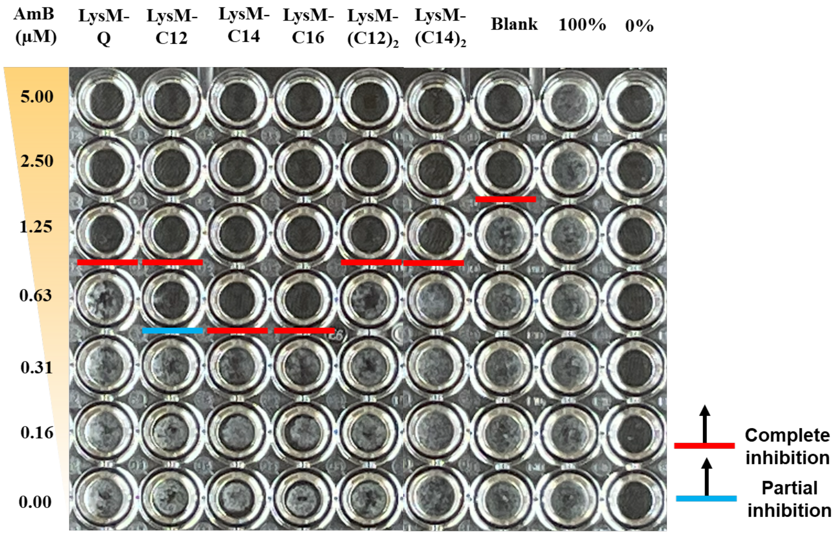


**Figure S4**. Representative image of a 96-well plate after culturing *T. viride* in the presence of 0−5 μM of AmB with 1 μM of each sample at 60h.

### 2-4. Results of DLS measurements

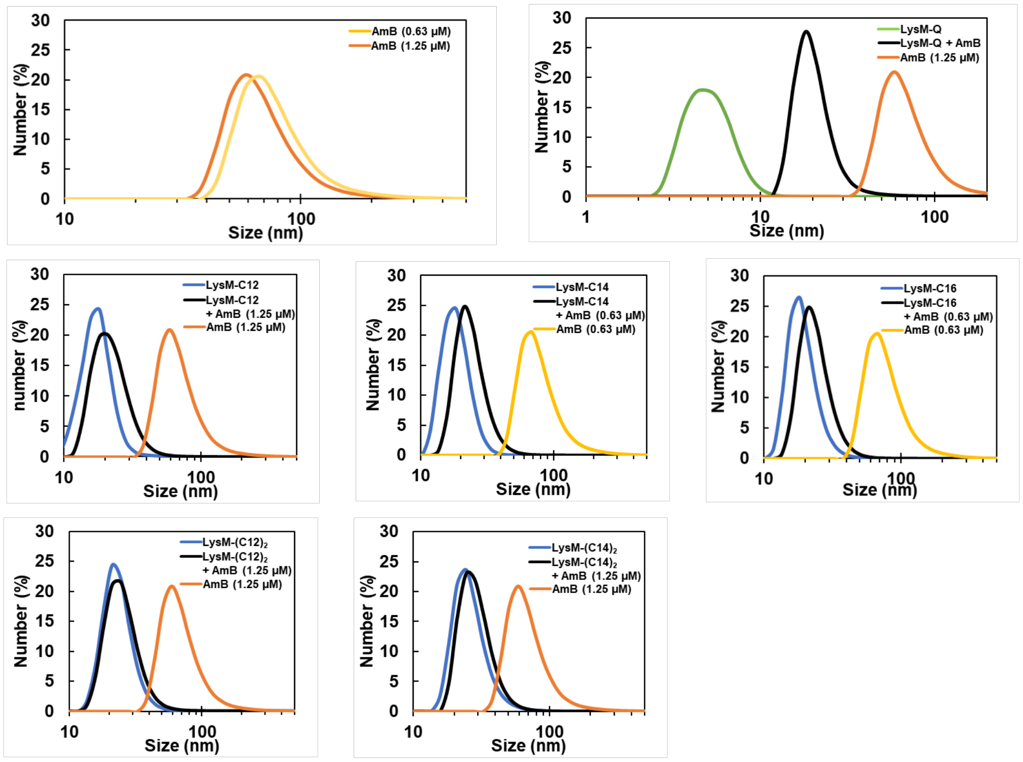


**Figure S5**. DLS measurements of AmB with LysM-Q or LysM-lipid in 20 mM NaPi, pH 7.4 at 25°C.

**2-5. CLSM analysis of interaction with *T. viride***

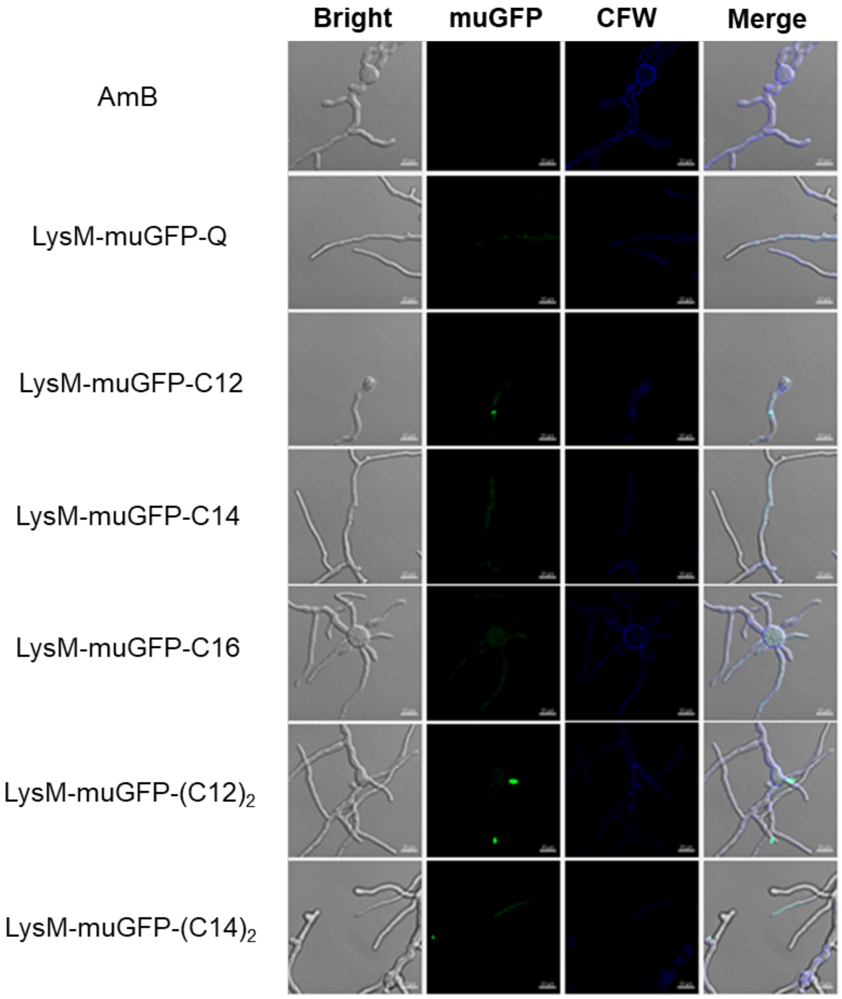


**Figure S6**. CLSM analyses of LysM-muGFP-Q or LysM-muGFP-lipid and combination without AMB in the presence of *T. viride* hyphae in 20 mM NaPi, pH 7.4, at 25°C (bars: 10 μm).

**2-6. CLSM analysis of LysM binding activity with α-chitin**

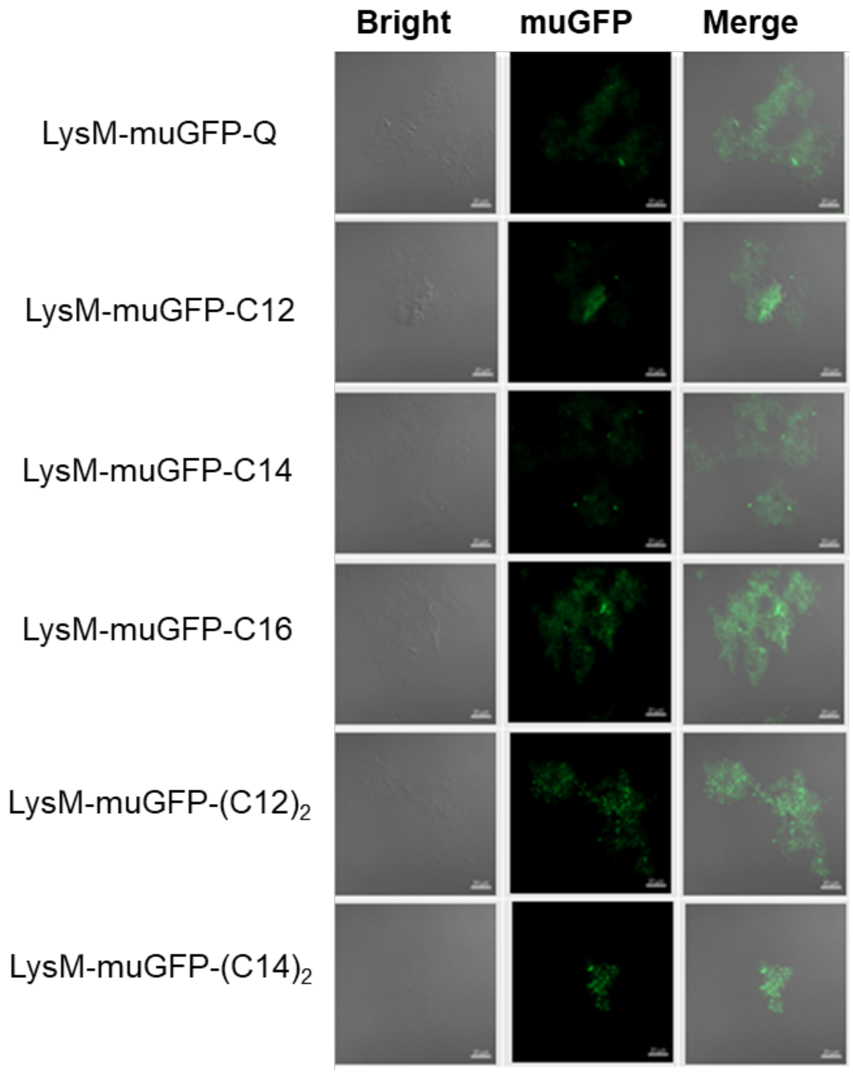


**Figure S7**. CLSM analyses of LysM-muGFP-Q or LysM-muGFP-lipid in the presence of 0.5% α-chitin in 20 mM NaPi, pH 7.4, at 25°C (bars: 10 μm).

#### 2-7. Cytotoxicity test of LysM-lipids with HeLa cells

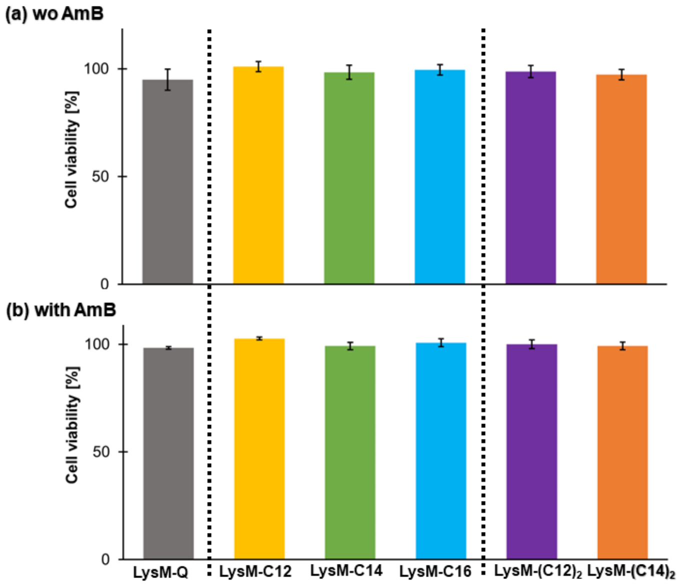


**Figure S8**. Cell viability of combination chitinase modified and AmB in HeLa cells (5,000 cell/well). Cell viability was quantified by Cell Counting Kit-8 (Dojindo) (A) without AmB and (B) with AmB.
